# Adopting Duplex Sequencing™ Technology for Genetic Toxicity Testing: A Proof-of-Concept Mutagenesis Experiment with N-Ethyl-N-Nitrosourea (ENU)-Exposed Rats

**DOI:** 10.1101/2023.05.08.539833

**Authors:** Stephanie L. Smith-Roe, Cheryl A. Hobbs, Victoria Hull, J. Todd Auman, Leslie Recio, Michael A. Streicker, Miriam V. Rivas, Gabriel A. Pratt, Fang Yin Lo, Jacob E. Higgins, Elizabeth K. Schmidt, Lindsey N. Williams, Daniela Nachmanson, Charles C. Valentine, Jesse J. Salk, Kristine L. Witt

**Affiliations:** Division of Translational Toxicology, NIEHS, Research Triangle Park, NC; Integrated Laboratory Systems, LLC (an Inotiv company), Research Triangle Park, NC; TwinStrand Biosciences, Inc., Seattle, WA

**Keywords:** DuplexSeq, Error-corrected NGS, ecNGS, Rat-50 Mutagenesis Assay, Transgenic rodent assay, *Pig-a* gene mutation assay

## Abstract

Duplex sequencing (DuplexSeq) is an error-corrected next-generation sequencing (ecNGS) method in which molecular barcodes informatically link PCR-copies back to their source DNA strands, enabling computational removal of errors by comparing grouped strand sequencing reads. The resulting background of less than one artifactual mutation per 10^7^ nucleotides allows for direct detection of somatic mutations. TwinStrand Biosciences, Inc. has developed a DuplexSeq-based mutagenesis assay to sample the rat genome, which can be applied to genetic toxicity testing. To evaluate this assay for early detection of mutagenesis, a time-course study was conducted using male Hsd:Sprague Dawley SD rats (3 per group) administered a single dose of 40 mg/kg N-ethyl-N-nitrosourea (ENU) via gavage, with mutation frequency (MF) and spectrum analyzed in stomach, bone marrow, blood, and liver tissues at 3 h, 24 h, 7 d, and 28 d post-exposure. Significant increases in MF were observed in ENU-exposed rats as early as 24 h for stomach (site of contact) and bone marrow (a highly proliferative tissue) and at 7 d for liver and blood. The canonical, mutational signature of ENU was established by 7 d post-exposure in all four tissues. Interlaboratory analysis of a subset of samples from different tissues and time points demonstrated remarkable reproducibility for both MF and spectrum. These results demonstrate that MF and spectrum can be evaluated successfully by directly sequencing targeted regions of DNA obtained from various tissues, a considerable advancement compared to currently used *in vivo* gene mutation assays.

**HIGHLIGHTS:**

- DuplexSeq is an ultra-accurate NGS technology that directly quantifies mutations
- ENU-dependent mutagenesis was detected 24 h post-exposure in proliferative tissues
- Multiple tissues exhibited the canonical ENU mutation spectrum 7 d after exposure
- Results obtained with DuplexSeq were highly concordant between laboratories
- The Rat-50 Mutagenesis Assay is promising for applications in genetic toxicology

## 1 INTRODUCTION

Genetic toxicity testing batteries used for regulatory submissions evaluate the mutagenic potential of chemicals, pharmaceuticals, and other substances for the purpose of hazard identification, as mutagenicity increases the risk for cancer, birth defects, and genetic disease. Standard testing batteries are designed to detect different kinds of heritable changes to DNA, including point mutations (changes in one or a few DNA base pairs, i.e., base substitutions, frameshifts, indels) and chromosomal damage (changes in chromosome structure or number) [1]. A longstanding goal in the field of genetic toxicology has been to evaluate endogenous and induced mutations in any tissue of any organism [2]. However, due to the inherent difficulty of detecting the extremely low frequency of point mutations induced by mutagens (i.e., in exposed tissues or cells that have not undergone natural cloning via tumor formation or reproduction of offspring), phenotypic selection assays, reflecting mutagenic events that occurred in protein-coding genes, emerged over the years as the most feasible approach to evaluate mutations *in vitro* and *in vivo*. These assays include bacterial reverse mutagenicity assays (a.k.a. the Ames test) [3] and several mammalian cell mutagenicity assays [4,5] for evaluation of gene mutations *in vitro* (e.g., *HPRT, XPRT, TK*), and the transgenic rodent assay (TGR) [6] for evaluation of gene mutations *in vivo*. During the past decade, the TGR assay was the only *in vivo* gene mutation assay that had an Organization for Economic Cooperation and Development (OECD) test guideline and is the closest the field has come to evaluating gene mutations in mammalian tissues [7]. The TGR assay relies on *ex vivo* phenotypic selection conferred by mutations in a single bacterial or viral gene, multiple tandem copies of which are, in nearly all cases, integrated into a single site of the rodent genome [8]. The *Pig-a* gene mutation assay, for which an OECD test guideline was recently adopted in 2022, is another option for evaluating *in vivo* mutagenesis [7]. Unlike phenotypic selection assays, the *Pig-a* assay directly detects a change in phenotype via loss of a cell surface marker that occurs due to inactivating mutations in the *Pig-a* gene of erythroid precursor cells, detected by high-throughput flow cytometric screening of millions of red blood cells [9]. Assessment of mutagenicity is limited to the hematopoietic compartment, and unlike TGR assays, mutation spectrum cannot be obtained.

Massively parallel next-generation sequencing (NGS) would appear to be a present-day solution to the challenge of evaluating point mutations *in vivo* or *in vitro*; however, due to artifacts that arise during sample preparation, amplification, and miscalls in the sequencing process, the ∼1×10^−3^ raw error rate for direct and high-throughput DNA sequencing is far greater than the estimated level of endogenous mutagenesis of ∼1×10^−8^ for mammalian cell DNA [10,11]. Duplex sequencing (DuplexSeq) is an error-corrected NGS (ecNGS) approach commercialized by TwinStrand Biosciences, Inc. that overcomes the poor sensitivity of standard NGS for detecting rare mutations [12]. DuplexSeq involves tagging short duplexes (∼300 bp) of fragmented DNA before strand separation and PCR amplification with a strategy that allows strand copies to be tracked back to their source strands informatically. A consensus-making algorithm is then used on all strand copies from a single source duplex to reduce errors and reconstruct the original molecule, revealing true mutations as complementary nucleotide changes at the same position on both strands. The resulting very low error rate of DuplexSeq, ∼1×10^−8^ – 1×10^−7^, enables direct detection of endogenous mutagenesis processes via measurements of mutation frequency (MF) and spectrum in any nucleated cell type of any organism, including humans.

The ability to directly sequence DNA for rare mutations has many applications, including the detection of point mutations for the purpose of genetic toxicity testing [13,14]. To increase the applicability of DuplexSeq for rodent mutagenesis studies, TwinStrand has created generalizable mutagenesis panels and assays for rats and mice that could supersede the use of specialized transgenic rodent models by directly sequencing DNA for mutations. The panels are designed to quantify MF and spectrum at a genomically representative set of 20 targets of ∼2.4 kb each (for a total of ∼50 kb) that are scattered across nearly all autosomes, covering both genic and intergenic regions. The panels are optimized for targets that are not challenging to sequence or align, such as those without excessive repetitive elements, lengthy homopolymers, closely related pseudogenes, and extremes of GC base pair content. The panels also exclude known cancer-related loci to avoid potential bias in MF due to positive or negative selective pressures. Recently, MF and spectrum obtained by the *cII* plaque assay using DNA from BigBlue® mice exposed to either N-ethyl-N-nitrosourea (ENU) or benzo[a]pyrene (B[a]P) were replicated by DuplexSeq [15]. A high level of concordance was also observed between DuplexSeq and mutagenesis of the *lacZ* transgene in the bone marrow of MutaMouse animals exposed to B[a]P [16] or when exposed to procarbazine [17]. An analogous human version of the DuplexSeq mutagenesis assay panel was also used successfully to identify ENU-induced mutations in human lymphoblastoid TK6 cells, an OECD-recommended cell line for genetic toxicity testing [18] and ethyl methanesulfonate (EMS)-induced mutations an *in vitro* air-liquid-interface airway tissue model used to evaluate the toxicity of inhaled substances [19].

This work explored the potential to adopt the TwinStrand DuplexSeq Rat-50 Mutagenesis Assay as an *in vivo* genetic toxicity test for mutation induction and mutation spectrum shift. Male Hsd:Sprague Dawley SD rats, typically used for toxicity testing, were administered a single dose of 40 mg/kg ENU, a well-characterized mutagen. MF and spectrum were evaluated in DNA isolated from stomach, bone marrow, blood, and liver tissues collected at 3 h, 24 h, 7 d, and 28 d after exposure. Rats exposed to vehicle were evaluated at 28 d after treatment. Tissues were chosen based on rapid versus slow rates of cell proliferation and site-of-contact as per the tissue selection recommendations of the TGR assay, which is typically conducted as a 28-day study [7]. Time points were chosen to model short-term toxicity testing and to evaluate how soon fixation of ENU mutations increases MF and shifts the endogenous spectrum to the known ENU mutational signature as detected by DuplexSeq. In alignment with the 3R principles (replacement, reduction, and refinement) for animal research [20], it was investigated whether 3 rats per group would be sufficient to reliably identify ENU-dependent mutagenesis using an ecNGS technology as sensitive as DuplexSeq. A subset of samples representative of different tissues and time points was analyzed to evaluate transferability of the technology and reproducibility of findings between laboratories. Finally, blood obtained from the same rats used for the DuplexSeq study was also analyzed using the *Pig-a* gene mutation assay at the 28-d time point (also the typical study duration time for the *Pig-a* assay) for a qualitative comparison of DuplexSeq (which is not restricted to evaluation of protein-coding genes) to a single-locus endogenous gene assay.

## 2 MATERIALS AND METHODS

### 2.1 Animal Treatment and Tissue Collection for Duplex Sequencing and *Pig-a* Gene Mutation Assay

All procedures complied with the Animal Welfare Act Regulations, 9 CFR 1-4, and animals were handled and treated according to the *Guide for the Care and Use of Laboratory Animals* [21]. Male Hsd:Sprague Dawley SD rats (Envigo Laboratories, Frederick, MD), 8 to 11 weeks of age, were housed 2 – 3 per polycarbonate cage with micro-isolator tops and absorbent heat-treated hardwood bedding (Northeastern Products Corp., Warrensburg, NY) and polycarbonate tubes for enrichment. Rats were provided certified Purina Pico Chow No. 5002 (Ralston Purina Co., St. Louis, MO) and reverse osmosis treated tap water *ad libitum*. Room temperature and humidity were approximately 20-25 °C and 30-70%, respectively. Lighting was on a 12/12-h light/dark cycle.

Rats were assigned to a dose group such that the mean body weight of each group was not statistically different from any other group using analysis of variance (ANOVA) (Statistical Analysis System version 9.2, SAS Institute, Cary, NC). Rats received a single oral gavage administration of freshly prepared 40 mg/kg N-ethyl-N-nitrosourea (ENU) (Sigma Aldrich, St. Louis, MO) dissolved in PBS (Nova-Tech, Kingwood, TX). Control rats received a single oral gavage administration of PBS (vehicle). Dose formulations were administered via oral gavage at a dose volume of 10.0 mL/kg body weight and dose volume was based on individual animal body weight.

Rats were euthanized by carbon dioxide asphyxiation and death was confirmed by exsanguination. Blood and tissues were harvested for duplex sequencing at 3 h, 24 h, 7 d, and 28 d post administration of ENU, and 28 d post administration of vehicle (3 rats per ENU time point and for the vehicle control), for duplex sequencing. Two frozen samples were prepared for each tissue type. Approximately 200 μL of whole blood obtained via cardiac puncture were collected into tubes, frozen on dry ice, and transferred to a -80 °C freezer. For the *Pig-a* gene mutation assay, additional blood (100 μL) was collected from vehicle-and ENU-treated rats at the 28-day timepoint, transferred to freezing solution (Litron Laboratories, Rochester, NY), and then to a -80 °C freezer according to the manufacturer’s instructions. Liver was rinsed in cold mincing buffer [Mg^2+^ and Ca^2+^ free Hanks Balanced Salt Solution (H6648-500ML, MilliporeSigma, St. Louis, MO), 10% v/v DMSO, and 20 mM EDTA pH 7.4-7.7], cut into 5 mm^3^ sections, placed into tubes, frozen on dry ice, and transferred to a -80 °C freezer. Bone marrow was flushed from the femur with 300 μL PBS (using an 18-gauge needle fitted onto a syringe) into microfuge tubes maintained on ice. Tubes were centrifuged at 1000 RPM for 5 min at 4 °C, the supernatant was aspirated, and the tubes containing the cell pellets were transferred to a -80 °C freezer. Stomach was cut open and washed using cold mincing buffer. Forestomach was discarded and the glandular stomach was placed in cold mincing solution and incubated on ice for 15 – 30 min. Surface epithelium was gently scraped twice using a Teflon scraper and discarded to remove surface contaminants. The gastric mucosa was rinsed with cold mincing buffer and the stomach epithelium was scraped 4 – 5 times in 500 μL mincing solution with the Teflon scraper to release cells. The mincing solution with the released epithelial cells was transferred to a microfuge tube, gently mixed, divided into two tubes, and centrifuged at 1000 RPM for 5 min at 4 °C; the supernatant was aspirated, and the tubes containing the cell pellets were transferred to a -80 °C freezer.

Due to the sensitivity of DuplexSeq, care was taken to avoid cross-contamination of samples during necropsy. Tools and necropsy stations were cleaned with 70% ethanol and RNAse*Zap*™ (ThermoFisher Scientific, CA) between each animal. To avoid contamination between organs, each organ was placed in its own clean, plastic petri dish and rinsed twice with cold mincing solution to remove blood. Tissues were then processed using disposable forceps. Each technician was designated to process only one type of organ. Tubes containing the processed samples were kept separate from each other until the end of necropsy when they were placed into boxes for transfer to a -80 °C freezer. Frozen tissue samples were shipped by FedEx overnight delivery to TwinStrand Biosciences, Inc., for DNA extraction and duplex sequencing.

#### DNA Extraction

DNA extractions were performed at TwinStrand Biosciences, Inc. with the Qiagen DNeasy Blood and Tissue Kit (69504; Qiagen) using the manufacturer’s protocol, with the exception that cell lysis was carried out at 37 °C for 10 min for blood and bone marrow, and 2 h for stomach and liver tissue. Extractions used approximately 200 total μL of whole blood (2 columns per sample with up to 100 μL blood per column, if available), 20 μL of bone marrow pellet, 5 mg of gastric mucosa pellet and 25 mg of liver tissue. DNA concentration was quantified using a Qubit™ instrument with high sensitivity reagents (Q32854; Thermo Scientific.)

#### Duplex Sequencing

For each sample, 375 ng (in some cases less when material was limiting) of extracted genomic DNA (gDNA) were fragmented to a median size of ∼300 base pairs using a Covaris ME220 ultrasonic shearing system (Covaris, Woburn, MA). For samples with a DNA integrity number (DIN) < 7 as assessed on an Agilent 2200 TapeStation (Agilent, Santa Clara, CA), 500 ng to 1 μg gDNA were fragmented if available. Paired-end Illumina sequencing libraries were created from fragmented gDNA using the DuplexSeq Rat-50 Mutagenesis Assay library preparation kit and protocol (06-1007-03 Rev. 1.0; TwinStrand Biosciences, Seattle, WA). The protocol (significantly optimized from Kennedy et al., 2014 [22]; carried out as described in Valentine et al., 2020 [15]) includes steps of end-repair, A-tailing, ligation of DuplexSeq adapters, and treatment with a conditioning enzyme cocktail to remove chemically damaged bases prior to PCR with unique dual index-containing primers. After template indexing and amplification, two tandem rounds of hybrid selection for mutagenesis target enrichment were carried out using a pool of biotinylated oligonucleotides, followed by washes and a final PCR step with P5/P7 primers. All final DuplexSeq libraries were quantified, pooled, and then sequenced using 150 bp paired-end reads targeting 200 million paired end reads per sample (100 million clusters) on an Illumina NovaSeq 6000 S4 flow cell using vendor-supplied reagents and version 1.0 chemistry.

#### Design of the Rat Mutagenesis Assay Panel

The DuplexSeq Rat-50 Mutagenesis Assay hybrid selection panel used in this study was designed and developed by TwinStrand Biosciences to capture contiguous territories of uniquely mapping regions of the rat genome to be used as genome-representative surrogates for genome-wide mutagenesis measurement. The panel was optimized to maximize hybrid selection capture uniformity and minimize potential alignment and variant calling technical artifacts. The panel is composed of twenty ∼2.4 kb contiguous baited regions for a total baited territory of ∼50 kb. The regions included are of a similar nucleotide composition, trinucleotide composition, %GC content, genic-to-intergenic ratio, and coding-to-non-coding ratio, as compared to the entire rat genome. Sites included in the panel have no obvious role in oncogenesis based on manual review of rat, human, and mouse homolog gene annotations (when available) to limit the potential for bias in mutagenesis measurements that could arise in genomic regions under positive or negative selection.

#### Bioinformatic Processing

Bioinformatic processing was carried out as per the methods described in Valentine et al. [15]. Alignment was performed using the BWA aligner version 0.7.15 [23] against the *Rattus norvegicus* reference genome rn6 (Rnor_6.0; GCF_000001895.5). Post-processing of the duplex consensus alignments included balanced overlap hard clipping and 5′ end-trimming to limit the generation of bias or technical artifacts during the subsequent variant calling process. Duplex consensus alignments were filtered to retain only those which were unambiguously from the *Rattus norvegicus* genome assembly using the taxonomic classifier Kraken v1 [24]. Somatic and germline variants were called using VarDictJava in tumor-only mode (v1.8.2) [25]. Loci with variant sub-clones present at outlier variant allele frequencies (VAF) in samples where other samples within the cohort had germline variant allele frequencies were omitted from downstream analysis, as these sites could be the result of intra-cohort contamination. Thirty-six such genomic single-base loci were omitted.

#### DuplexSeq Statistical Analysis

Regions within 10 bp upstream or downstream of any detected germline indels were excluded from the analysis due to the potential for alignment-based artifacts. MF was calculated as the number of unique somatic variant alleles (*e*.*g*., single nucleotide variants, multi-nucleotide variants, insertions, deletions, and structural variants) divided by the number of non-ambiguous duplex consensus bases across the entire panel territory for each sample. For example, if a specific type of mutation at a specific genomic site in the panel was found in three independent duplex consensus alignments from the same sample, the variant would only be counted once. This method of calculating MF limits the potential for jackpot events to bias MF quantitation should a substantial clonal expansion of a cell lineage carrying the mutation have occurred, either postnatally or as a result of embryonic somatic mosaicism. Simple base substitution spectra were generated from the count of somatic base substitutions rectified into pyrimidine-space and then normalized by the frequencies of reference nucleotides in the panel territory. Trinucleotide base substitution spectra were generated in the same fashion as simple base substitution spectra except that final normalization was performed using the number of 3mers in the panel territory’s reference sequence. Linear regression was run with the “lm” function in R (v3.6.3) where MF was set as the response variable and predictor variables were treatment, tissue, and time of exposure. Variation of MF within each treatment group was evaluated with the coefficient of variation (CoV).

#### Analysis of Mutation Frequency by Target

Each unique somatic variant allele was grouped by the panel target it overlapped with to evaluate MF for each of the twenty target regions. MF for a treatment group was calculated as the number of unique somatic variant calls in a target region divided by the number of non-ambiguous duplex consensus bases across the same target region for all animals within a treatment group.

#### Interlaboratory Reproducibility Study: DNA Extraction at Inotiv

DNA extractions on duplicate tissue samples of blood, gastric mucosa, and liver were performed at ILS, LLC with the Qiagen DNeasy Blood and Tissue Kit using the manufacturer’s protocol. Extractions used approximately 50 μL of whole blood, 5 mg of gastric mucosa pellet, and 25 mg of liver tissue. A blood sample was homogenized with a handheld pestle motor mixer (Argos, Thomas Scientific, NJ) if it appeared to be clotted. Cell lysis was carried out at 37 °C for 10 min for blood and 2 h for stomach and liver tissue. For stomach and liver samples that were not completely lysed after 2 h, an additional 20 μL of Proteinase K were added, followed by overnight incubation at 37 °C.

#### Interlaboratory Reproducibility Study: Duplex Sequencing at Inotiv

For each sample in the interlaboratory comparison, 100-500 ng of purified DNA were fragmented in TE-low buffer (10mM Tris HCl, 0.1 mM EDTA) using a Covaris ME220 ultrasonic shearing system to a median size of ∼300 bp as determined by the Agilent 2100 High Sensitivity Assay (Agilent, Santa Clara, CA). Libraries were generated using the DuplexSeq Rat-50 Mutagenesis Assay library preparation kit and protocol using version 1.0 chemistry as described above. All final DuplexSeq libraries were quantified, pooled in batches of 4 libraries, and then sequenced with 150 bp paired-end reads (targeting 200 M total reads, 100 M in each direction) on an Illumina NextSeq 500 High Output flow cell using vendor supplied reagents and version 2.5 chemistry. Demultiplexed sequence files were merged using the FASTQ Toolkit (version 2.25; https://basespace.illumina.com/apps/4457453/FASTQ-Toolkit?preferredversion) and uploaded to the TwinStrand portal on DNAnexus for mutagenesis analysis (https://platform.dnanexus.com/app/twinstrandbio-mutagenesis).

#### *Pig-a* Gene Mutation Assay

On the day of analysis, frozen blood was washed out of freezing solution (MutaFlow^®^ Rat Blood Freezing Kit, Litron Laboratories, Rochester, NY), and labeling and flow cytometric analyses of reticulocytes (RETs) and red blood cells (RBCs) were performed using a MutaFlow^®^ kit (Litron Laboratories, Rochester, NY) and a Becton-Dickinson FACSCanto™ II flow cytometer (Sunnyvale, CA). The % RETs and frequencies of *Pig-a* mutant phenotype RETs and RBCs were calculated based on pre- and post-column analyses [26].

## 3 RESULTS

DuplexSeq yield for all 59 samples was high with each sample generating at least 50 million non-ambiguous duplex bases (mean = 422 million, range of 63 million to 1,212 million), and hybrid selection was robust, with all but one sample having at least 85% of the duplex bases residing on-target (Supplementary Data Fig. 1). Samples were assessed for germline variant sequences unique to each animal (fingerprinting) and then for potential low-level contamination with DNA from organisms other than rats. A fingerprint analysis revealed that only two, not three, independent samples for the ENU 24 h bone marrow exposure group were submitted to TwinStrand for library preparation (data not shown) reducing the total count of unique samples analyzed from 60 to 59. Six samples from different tissues and time points among the 59 samples that were analyzed showed very low levels (< 0.01% of reads) of contamination with human DNA (Supplementary Data Table 1). Sequences of human origin were removed before analysis of MF and spectrum.

The MF for each sample over time is shown in Fig. 1A-D. Supplementary Data Tables 2 – 5 report the MF for each data point in Fig. 1A-D, the mean MF for each exposure group, and 95% Wilson confidence intervals, along with the number of unique mutations and mean duplex depth per sample. The mean MF for vehicle control tissues ranged from 0.47×10^−7^ for stomach to 1.7×10^−7^ for blood. Significant increases in MF were detected 24 h after exposure to ENU for stomach and bone marrow tissues. At 7 d after exposure to ENU, significant increases in MF were detected in all four tissues. For stomach, bone marrow, and blood tissues, increases in ENU-dependent MF were roughly linear up to 7 d, at which point they appeared to plateau through 28 d after exposure, whereas increases in MF were linear in liver tissue up to 28 d after exposure. At 28 d, MF was induced 40-, 8-, 8-, and 5-fold in stomach, bone marrow, blood, and liver tissue, respectively. A CoV analysis showed that in some cases, especially for samples from ENU-exposed rats, there was very little variation in MF among biological replicates; stomach tissue at the 3 h and 24 h time points were notable exceptions, potentially due to the inherently more complex and variable nature of scraping the gastric mucosa to release epithelial cells (Supplementary Data Fig. 2, Supplementary Data Table 6). A limitation is that with three samples per time point and tissue, one different value can considerably skew the CoV analysis.

**Fig. 1A – D.**
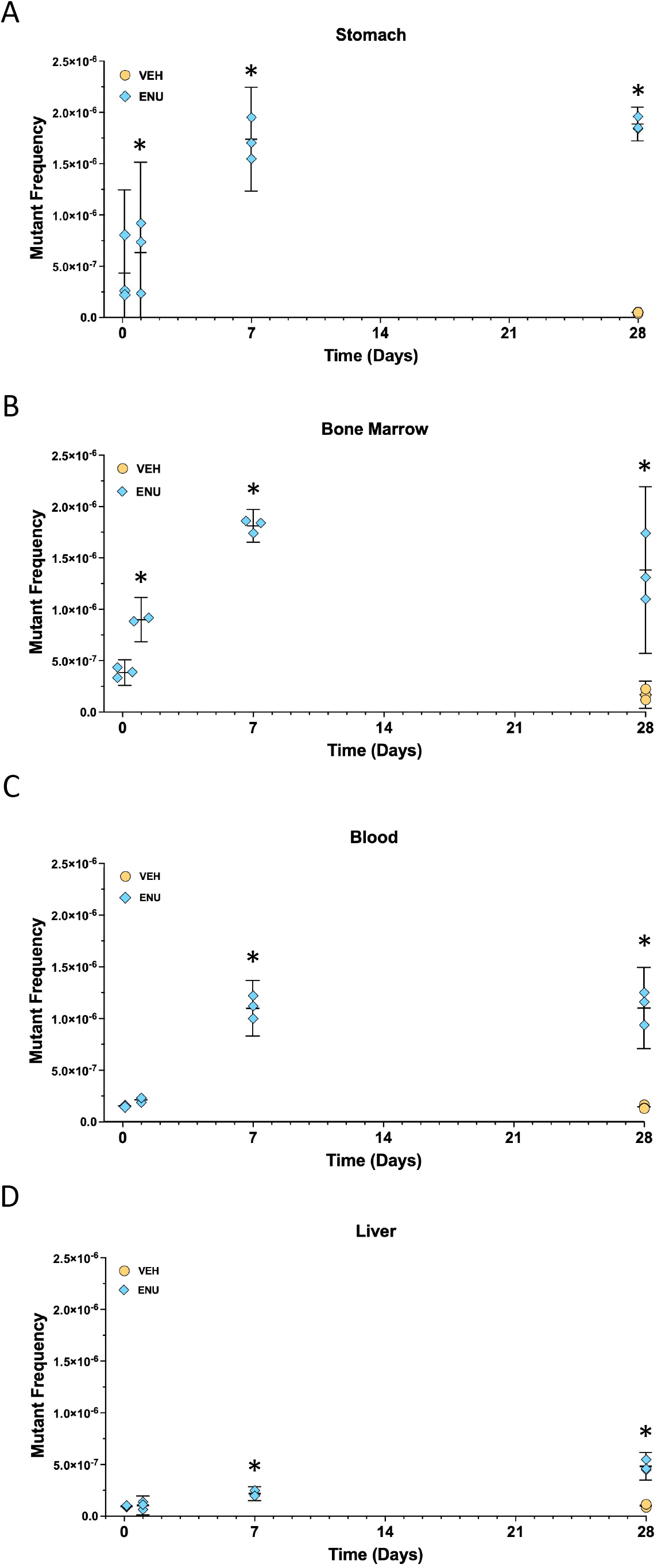
Mutation frequency over time for stomach (A), bone marrow (B), blood (C), and liver (D) tissue samples at exposure timepoints 3 h, 24 h, 7 d, and 28 h. Note that vehicle control rat tissues were sampled only at 28 d. Horizontal bar indicates the mean, vertical bars indicate lower and upper 95% confidence intervals. ^*^P < 0.05.

To investigate whether the mutagenic effects of ENU varied at each target, the MF for each treatment group at each time point was determined for each of the 20 targets for stomach (2A), bone marrow (2B), blood (2C), and liver (2D). The targets for stomach and liver were ordered based on the target with the highest MF at the 28-day time point (left most) to the region with the lowest MF at the 28-day time point. The targets for bone marrow and blood were ordered based on the target with the highest MF at the 7-day time point (left most) to the region with the lowest MF at the 7-day time point. A comparison of MF at each target for each tissue at the 28-d time point, by which time overall MF plateaued or was highest for each tissue, is shown in Fig. 3. The targets for each tissue were ordered based on the region with the highest MF in stomach (left most) to the region with the lowest MF in stomach. Although this study was not sufficiently powered to assess whether there were significant differences in MF across the different targets, the following observations can be made. In all four tissues, all 20 targets showed increases in MF over time after exposure to a single dose of ENU. For each tissue, the relative increases in MF at each target and at each time point were largely similar (Fig. 2A-D). In Fig. 3, the highest response versus the lowest response for a target differed by 3.8-fold, 3.0-fold, 2.6-fold or 2.4-fold for stomach, bone marrow, blood, and liver, respectively. Also, while all 20 targets were responsive to ENU exposure, MFs tended to be lower for the intergenic targets on chromosomes 7, 12, and 19, and the genic target on chromosome 16, at 28 d after exposure; however, across all time points, no single target region was consistently lower, or higher, than others.

**Fig. 2A – D.**
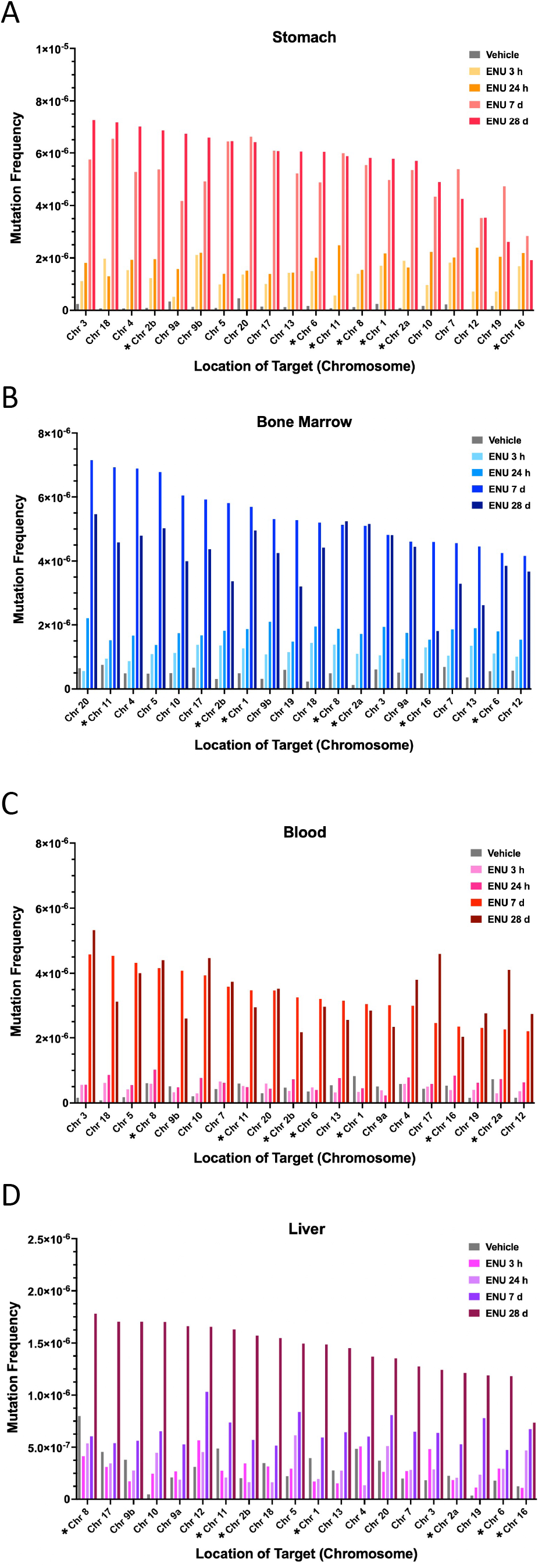
Mutation frequency at each target in the DuplexSeq Rat-50 Mutagenesis Assay for stomach (A), bone marrow (B), blood (C), and liver (D) tissue sampled. ^*^Genic targets.

**Fig. 3.**
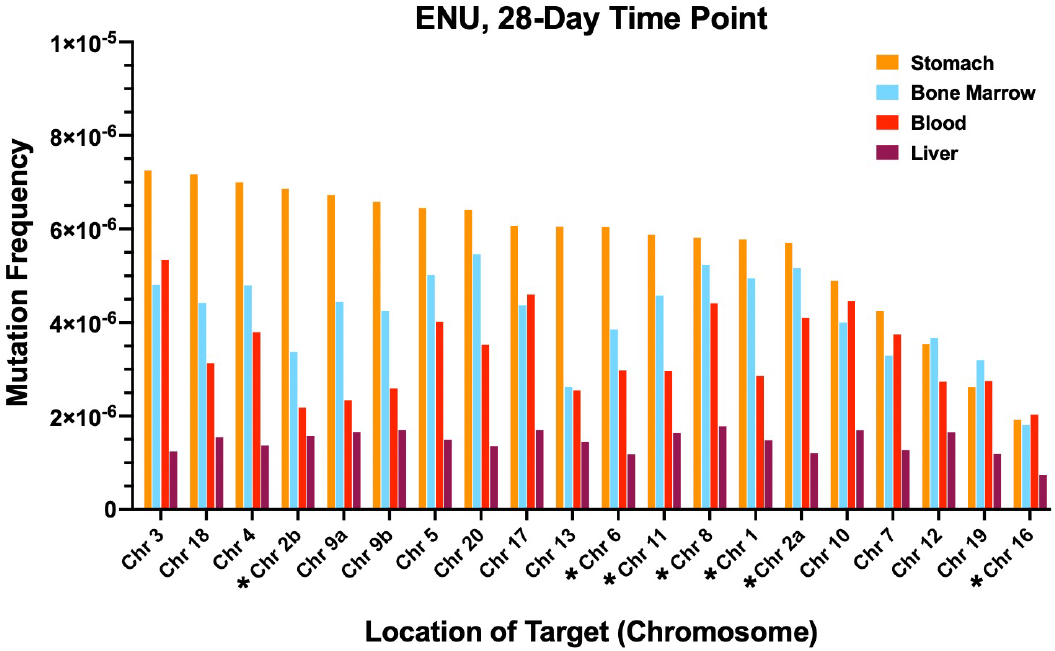
Mutation frequency at each target in the DuplexSeq Rat-50 Mutagenesis Assay for each tissue at the 28-d time point. ^*^Genic targets.

Base substitution count and proportion are shown for each sample in Fig. 4. The simple base substitution mutation spectra were similar for all four tissues obtained from vehicle control animals. The vehicle control mutation spectra were dominated by C:G > A:T and C:G > G:C transversion mutations associated with endogenous oxidative damage to guanine bases, and C:G > T:A transitions, likely resulting from unrepaired spontaneous deamination events at cytosine and 5-methyl-cytosine bases. Variability among the vehicle control samples was mostly due to statistical factors of under-sampling, as indicated by relatively low mutation counts per sample. At 3 h and 24 h after exposure to ENU, the simple mutation spectra for all four tissues were dominated by C:G > T:A transitions. At 7 d after exposure to ENU, the proportion of mutation subtypes known to be associated with ENU, T > C transitions and T > A transversions [27,28], increased considerably compared to earlier time points. T:A > C:G transitions comprised approximately 29%, 30%, 32%, and 40% of mutations for blood, bone marrow, stomach, and liver tissues, respectively, and T:A > A:T transversions made up approximately 19%, 26%, 26%, and 30% of those in liver, blood, bone marrow, and stomach tissues, respectively. T:A > G:C transversions, which are also associated with ENU exposure [29], increased at 7 d compared to earlier time points, and accounted for ∼6 – 8% of mutations across tissues. Taken together, ∼60 – 70% of the simple mutation spectra were associated with ENU exposure for each tissue at the 7-d time point. For all tissues, the ENU-dependent mutation spectra at 28 d were similar to those observed at 7 d after exposure. As with the MF, biological replicates generally showed little variation in mutation spectra across all tissues, except for the 3 h and 24 h stomach samples, which showed a high degree of variability for C > T mutations (Supplementary Data Fig. 3).

**Fig. 4.**
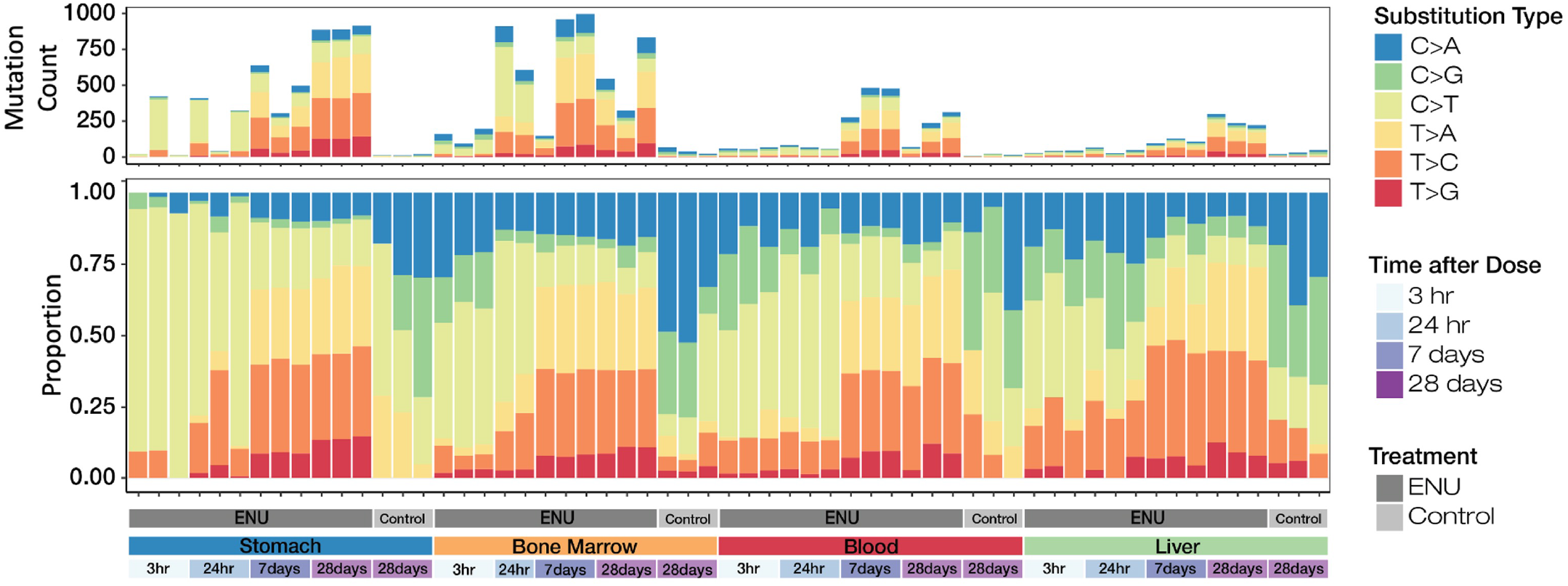
Unique mutation counts and normalized proportions of the 6 canonical base substitution types for each sample. Note that vehicle control rat tissues were sampled only at 28 d.

To evaluate the transferability of DuplexSeq technology and the reproducibility of data obtained using the Rat-50 assay, a subset of duplicate tissue samples underwent DNA extraction, library preparation, sequencing, and analysis using the TwinStrand DNAnexus cloud pipeline by ILS, LLC. Quality control parameters are shown for these samples in Supplementary Data Fig. 4, and a comparison of MF for samples processed by TwinStrand versus ILS is shown in Fig. 5A, B and Supplementary Data Table 7. The MFs obtained by the two laboratories, shown in Fig. 5A, had a very high degree of correlation (Pearson *r* = 0.986) (Fig. 5B). Furthermore, trinucleotide spectra developed from vehicle control or ENU-exposed stomach, blood, and liver samples processed at TwinStrand or ILS were highly concordant in a cosine similarity analysis (Fig. 6). Comparisons of trinucleotide spectra by proportion as obtained by TwinStrand versus ILS for stomach, blood, and liver samples are shown in Supplementary Data Figs. 5,6, and 7, respectively. Lastly, the pyrimidine-normalized trinucleotide (3-mer) counts in the DuplexSeq ™ Rat-50 mutagenesis panel (which excludes XY chromosomes) are proportional to the autosomal content of the *Rattus norvegicus* genome, Pearson r = 0.98 (Supplementary Data, Fig. 8).

**Fig. 5A, B.**
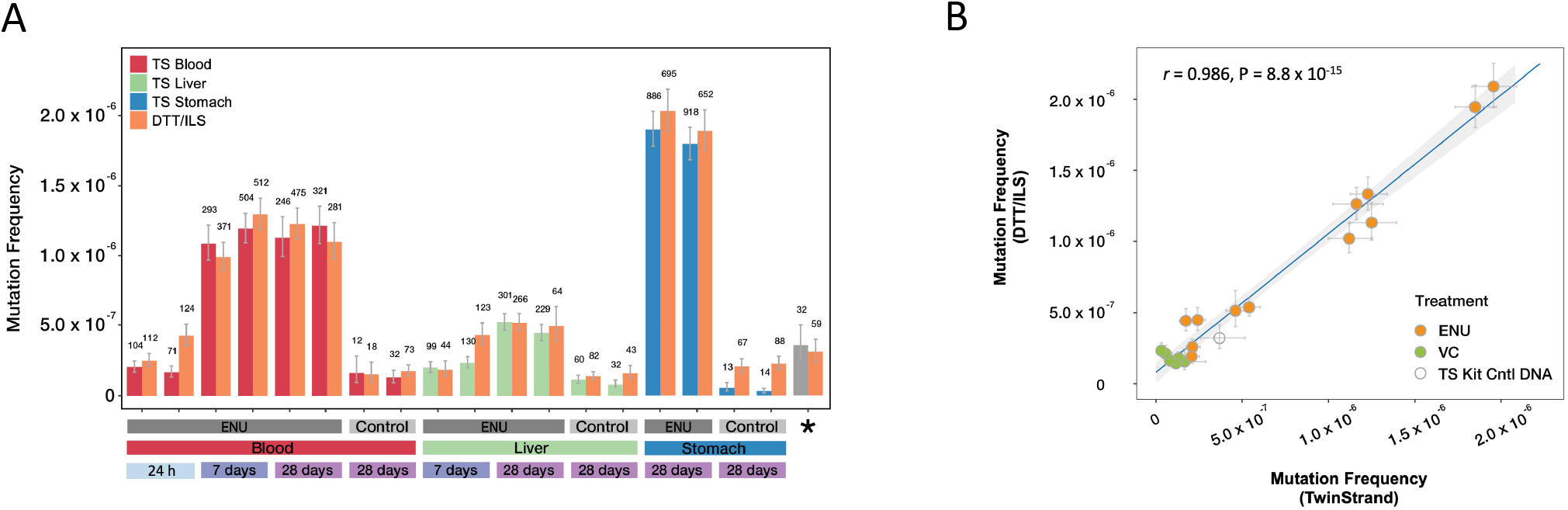
Comparison of mutation frequency for a subset of samples that underwent DNA extraction, library preparation, and sequencing at either TwinStrand Biosciences, Inc., or ILS, LLC. Error bars indicate lower and upper 95% Wilson confidence intervals. The number of unique mutations detected per sample is noted above each bar; ^*^ indicates a control male rat liver DNA provided in the DuplexSeq Rat-50 Mutagenesis Assay library preparation kit (A). Correlation analysis of TwinStrand versus DTT/ILS mutation frequencies shown in 5A; shaded area represents the 95% confidence interval of the regression line (B).

**Fig. 6.**
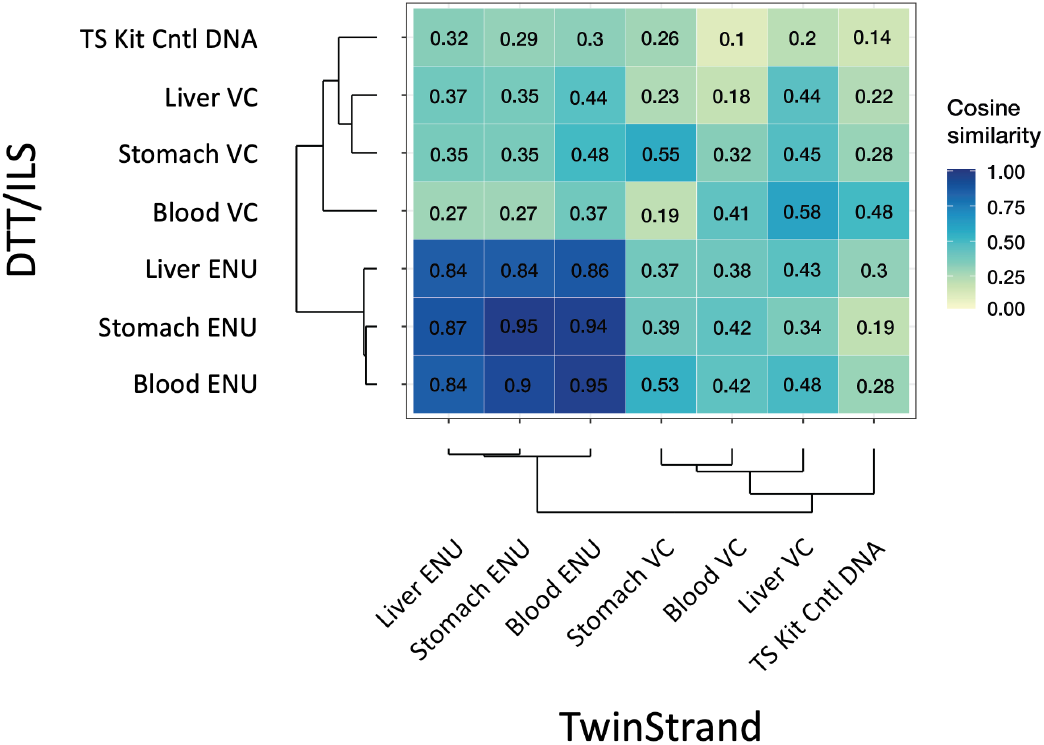
Cosine similarity analysis of trinucleotide sequence spectra obtained by TwinStrand Biosciences, Inc. versus ILS, LLC for the samples shown in Fig. 5A and Fig. 5B.

The same rats that were sampled for DuplexSeq analyses were also sampled for the *Pig-a* gene mutation assay at 28 d after exposure to ENU or vehicle control. A single exposure to ENU dramatically increased MF at the *Pig-a* locus in precursor erythroid cells (Table 1). The mutant RET frequency increased from 2.4×10^−6^ to 202.7×10^−6^ (84-fold over vehicle control), and the mutant RBC frequency increased from 0.9×10^−6^ to 107.3×10^−6^ (119-fold over vehicle control). The mutant RET and RBC vehicle control frequencies were within the historical vehicle control range for ILS.

**Table 1:**
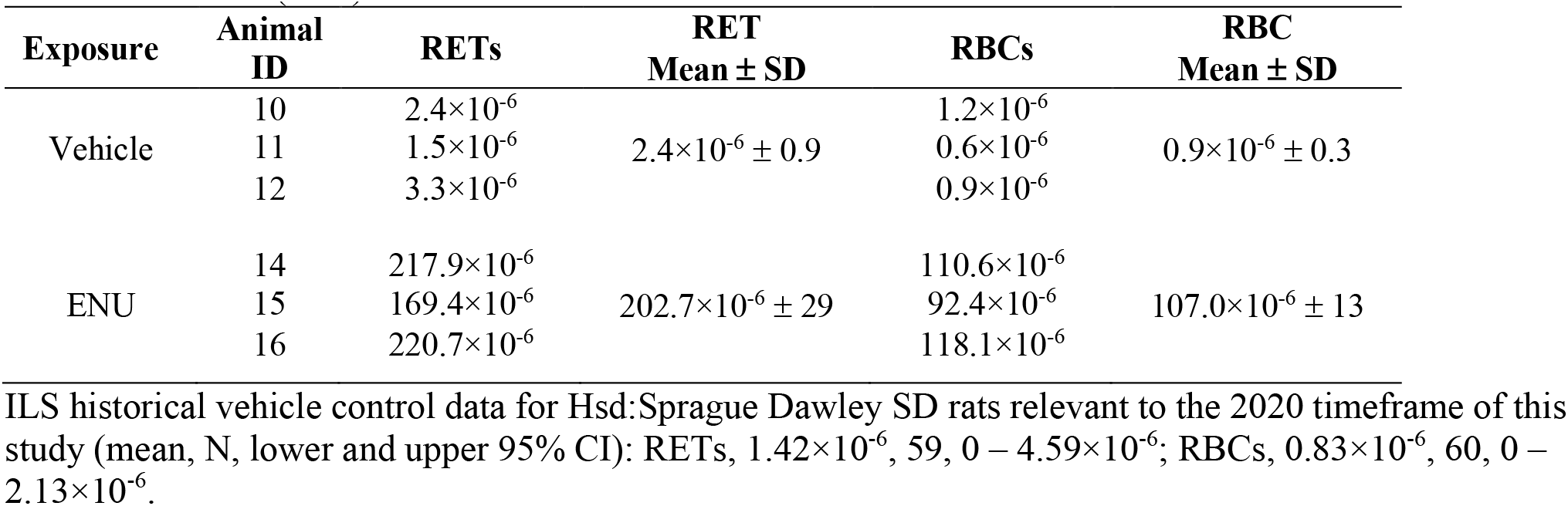
Mutant reticulocyte (RET) frequency and mutant red blood cell (RBC) frequency obtained from the *Pig-a* gene mutation assay at 28 days post-exposure to 40 mg/kg ENU or vehicle control (PBS)

| Exposure | Animal ID | RETs | RET Mean ± SD | RBCs | RBC Mean ± SD |
| --- | --- | --- | --- | --- | --- |
| Vehicle | 10 | $2.4 \times 10^{-6}$ | | $1.2 \times 10^{-6}$ | |
| | 11 | $1.5 \times 10^{-6}$ | $2.4 \times 10^{-6} \pm 0.9$ | $0.6 \times 10^{-6}$ | $0.9 \times 10^{-6} \pm 0.3$ |
| | 12 | $3.3 \times 10^{-6}$ | | $0.9 \times 10^{-6}$ | |
| ENU | 14 | $217.9 \times 10^{-6}$ | | $110.6 \times 10^{-6}$ | |
| | 15 | $169.4 \times 10^{-6}$ | $202.7 \times 10^{-6} \pm 29$ | $92.4 \times 10^{-6}$ | $107.0 \times 10^{-6} \pm 13$ |
| | 16 | $220.7 \times 10^{-6}$ | | $118.1 \times 10^{-6}$ | |

ILS historical vehicle control data for Hsd:Sprague Dawley SD rats relevant to the 2020 timeframe of this study (mean, N, lower and upper 95% CI): RETs, $1.42 \times 10^{-6}$ , 59, 0 – $4.59 \times 10^{-6}$ ; RBCs, $0.83 \times 10^{-6}$ , 60, 0 – $2.13 \times 10^{-6}$ .

## 4 DISCUSSION

Due to the technical challenges of identifying rare mutations, detection of point mutations *in vitro* and *in vivo* for the purpose of genetic toxicity testing has been limited to assays that evaluate phenotypic changes due mutations in protein-coding genes, mostly using biological selection-based techniques. The DuplexSeq strategy of comparing informatically linked complementary strands of DNA to distinguish true mutations from errors enables the use of NGS to directly evaluate mutations across the genome [12,14]. This work investigated the utility of the DuplexSeq Rat-50 Mutagenesis Assay designed by TwinStrand Biosciences, Inc. This platform uses a bait-and-capture approach to sample twenty representative ∼2.4 kb targets distributed across 18 of the 20 rat autosomes to detect mutations in several tissues typically evaluated in transgenic rodent mutation studies [7]; blood was also sampled to provide qualitative comparison to the *Pig-a* gene mutation assay.

In this time-course study, rats were administered a single dose of ENU, a mutagen that is absorbed and distributed throughout the body, and that reacts directly with DNA, produces a distinct mutational spectrum for which the mechanism-of-action has been characterized, and is commonly used as a positive control for the TGR assay [7,30]. Significant ENU-dependent increases in MF in stomach, bone marrow, blood, and liver tissue were detected at 7 d after exposure. For the rapidly proliferating stomach and bone marrow tissues, significant increases in MF were detected as early as 24 h after exposure. MF did not change appreciably between the 7- and 28-d time points for stomach, bone marrow, and blood tissues. Mutations continued to accumulate up to the 28-d time point in liver, a tissue that has slower cellular turnover compared to stomach, bone marrow, and blood. The lower level of mutagenesis in liver tissue compared to the other tissues examined in this study may be due to a combination of slower cellular turnover and a greater capacity for removing pro-mutagenic alkylation damage prior to fixation [8,31,32]. The liver also has comparatively high concentrations of glutathione, which detoxifies nitroso compounds [33,34]. This proof-of-principle experiment, albeit with a powerful mutagen, suggests that DuplexSeq could be used to obtain MF for the purpose of hazard identification from short-term toxicity studies, in alignment with 3R principles [20].

With ENU exposure, a clear change in mutation spectrum compared to the vehicle control group preceded significant increases in MF in all tissues. For stomach and bone marrow tissue, an increase in the C:G > T:A mutational fraction was apparent at 3 h (as well as 24 h), preceding significant increases in overall MF that first occurred at 24 h; for blood and liver tissue, an increase in this same mutational fraction was apparent at 24 h (as well as 3 h) preceding significant increases in overall MF that first occurred at 7 d. This observation highlights the sensitivity of DuplexSeq, such that changes in mutation spectrum can be detected before the occurrence of significant changes in MF, which are “buffered” to some degree by the larger baseline abundance of age-associated oxidation and deamination-mediated mutations. A change in ENU-dependent mutation spectrum over time was also observed, from mutagenic events arising from alkylation of guanine residues to those arising from alkylation of thymine residues. The C:G > T:A transition, observed at 3 and 24 h, is not considered to be a canonical mutation type for ENU; however, C:G > T:A transitions are the defining characteristic of ENU exposure in human lymphoblastoid TK6 cells, which lack O^6^-alkylguanine-DNA alkyltransferase (AGT) activity [35], and recently, the DuplexSeq Human-50 Mutagenesis Assay also identified significant increases in the proportion of C:G > T:A transitions in TK6 cells exposed to ENU [18]. AGT is a direct-reversal enzyme that removes O^6^-alkylation damage to DNA via covalent transfer to a cysteine residue, causing inactivation of the enzyme [36]. Because AGT is transcriptionally induced in response to DNA damage, including damage from ENU, we speculate that the early C > T transition signature of ENU in the rat tissues that were examined may have been due to constitutive levels of AGT that were insufficient to adequately repair O^6^-alkylation damage from a high dose of ENU. Although little is known about the dynamics of *in vivo* induction of AGT in response to ENU [37,38], one study showed that AGT enzymatic activity in rat liver tissue increased linearly from 1 to 3 days in response to a single dose of 2-acetylaminofluorine, and had decreased by 6 d [39]. In our study, for all 4 tissues, the mutation spectrum was dominated by base substitutions that are typically associated with ENU exposure, T > C transitions and T > A transversions, by 7 and 28 d [30,40-43]. These data indicate that the selection of time point may influence the conclusions to be drawn from mutational analysis of tissues when using a technology as sensitive as DuplexSeq.

DuplexSeq achieves high sensitivity and specificity compared to conventional NGS approaches by using sequencing information from informatically linked DNA strands to identify DNA mutations at a frequency as low as 1 in 100 million bases [14]. This sensitivity to detect variants is dependent on the average depth of sequencing and breadth of targeted regions sequenced (in this study, ∼50,000 bp). In this study, 50 samples had > 200 million informative duplex bases and all but 5 had at least 100 million (Supplementary Data Fig. 1) which enables comfortable quantification of MFs of ∼1×10^−7^ and greater (10 mutations per 100 million duplex base pairs). The deeper the sequencing or the greater the size of the targeted panel, the greater the number of informative bases obtained and the better the ability to resolve more subtle differences between the MF of two samples.

In the future, it will be necessary to develop a better understanding of the underlying spontaneous mutation rate for each tissue as detected by DuplexSeq, which is dependent on the frequency of cell division in each tissue and the total number of cell divisions that have occurred by the time of sample collection (*i*.*e*., the age of the animal). The current experiment using 3 animals per group showed a range of variability in the MF for each animal for a set of samples. The differences in MF between animals is most likely due to a combination of biological and technical factors. Possible biological factors include variations in response to a given chemical and differential cellular states that can influence a cell’s ability to repair damaged DNA. Technical factors most likely impacting this data set relate to variability inherent in certain sampling procedures requiring more handling (scraping of gastric mucosa) but could also include variables associated with manipulation of DNA or the DuplexSeq assay itself, in addition to the stochastic nature of probabilistic sampling at very low frequencies where mutation counts are low. In particular, the small numbers of mutations detected in vehicle control animals are underpowered to detect meaningful differences among these animals, even if they were to exist. Additional experiments are needed to obtain a better estimate of the variability in MF in animals and different tissues to develop best practice guidelines for the number of animals needed per experimental group for robust and reproducible results.

The amount of sequencing required to achieve a certain number of informative duplex bases is an important factor to consider for balancing sensitivity and expense when performing a DuplexSeq Mutagenesis Assay. In general, doubling the amount of DNA inputted into the assay will double the number of informative bases produced (and therefore the statistical confidence around a specific measured frequency) but it will also double the sequencing cost. While a relatively modest number of duplex bases is sufficient to detect a change in MF resulting from exposure to a strong mutagen in a very sensitive tissue after ample mutation fixation time (for example, ENU in stomach harvested at 7 days after exposure, which yielded an extreme 40-fold MF increase in this study), the sequencing requirements to robustly detect the more subtle changes in less sensitive tissues and at earlier time points or lower doses will be greater. For the greatest induction seen in this study, perhaps as little as 1/10^th^ of the data generated from those samples would have been sufficient to detect this induction with statistical confidence, but for a much weaker mutagen or lower dose expected to yield only a 50% increase in MF, a larger number of duplex bases would be needed. Significant future investigations across a broader range of mutagens and doses will be needed to determine the optimal amount of data for different applications.

The reproducibility study provides insights into the technical variability in the DuplexSeq results. There was a very high technical correlation in MF observed from the same samples that were processed in two different laboratories. The internally controlled ratiometric nature of the assay (mutations identified per bases sequenced) helps inherently normalize for differences in data yield across different sites or users. Whereas some significant differences in MF were observed among the cohort of samples processed at TwinStrand, the inter-animal variation (i.e., due to biology, dosing, or sampling variability) was drastically greater than the inter-laboratory variation (technical variability). Further experiments are warranted to assess the sources and frequency of limited inter-laboratory variability (for example, the 28-day stomach control samples in this study) to better understand the transferability and long-term reliability of DuplexSeq. It should also be noted that the very high sensitivity of DuplexSeq improves the reliability of sequencing data by detecting contamination of foreign DNA and, in the case of animal studies, inadvertent sample swaps by using germline mutational fingerprints as identifiers for samples.

An important consideration for any new technology is how it compares to established ones. In blood samples taken from the same animals and assessed at 28 d post exposure to ENU, the *Pig-a* assay detected 84-fold and 119-fold increases in mutant RET and mutant RBC frequencies, respectively, whereas the DuplexSeq Rat-50 Mutagenesis Assay detected 7.8-fold and 8.1-fold increases in MF for blood and bone marrow, respectively. The *Pig-a* assay appears to be much more sensitive for the detection of ENU-dependent mutagenesis compared to DuplexSeq. However, comparison of results obtained with these assays is limited for several reasons. First, the approaches are very different, as the *Pig-a* assay detects loss of the CD59 cell surface marker due to mutagenesis at the locus that encodes phosphatidylinositol glycan anchor biosynthesis class A, the protein needed for CD59 expression, versus measuring mutations identified per bases sequenced across different sites of the genome. Second, whereas *Pig-a* mutant frequencies reflect mutagenesis that occurred in erythroblasts, a variety of cell types are sampled when DNA is extracted from bone marrow, and sequencing DNA from blood detects mutations that occurred in leukocytes. Furthermore, these various cell types may differ in their sensitivity to the cytotoxic effects of ENU. Lastly, we speculate that the difference in mutational response to ENU might also be partly due to how DuplexSeq corrects for clonal expansions. DuplexSeq mitigates MF inflation from clonal amplification by counting unique mutations only once regardless of how many copies of that mutation are observed, limiting the effect of “jackpot” events. A closer comparison of these assays would require duplex sequencing of the *Pig-a* gene in isolated erythroblasts, as well as testing with mutagens that are not as strong as ENU. Despite these differences, the results are concordant between the two assays. Notably, whereas the *Pig-a* assay requires ≥ 14 d for detectable expression of the mutant phenotype, statistically significant ENU-dependent mutagenesis could be detected using DuplexSeq as early as 24 h after exposure of the hematopoietic compartment, with the advantage of characterizing the mutational signature of exposure.

This proof-of-principle study demonstrated that strategic sampling of the genome using DuplexSeq, developed as the DuplexSeq Rat-50 Mutagenesis Assay, is effective for quantitatively measuring chemically induced mutagenesis in rat tissues. Furthermore, results were highly reproducible between two laboratories independently carrying out the assay. How this technology will perform for various types of mutagens still needs to be determined, and a general understanding of the background spontaneous mutation rate and mutation spectra in untreated animals will be necessary to distinguish the DNA mutations associated with chemical exposure via MF or spectra. These findings and those reported by others [15-17] support research to explore the adoption of DuplexSeq for the purpose of creating standardized rodent genetic toxicity tests to meet regulatory needs for the detection of point mutations [13], similar to efforts made to develop OECD test guidelines for the TGR and *Pig-a* gene mutation assays. Compared to currently used assays, direct detection of mutations and identification of mutational spectra in animal models has greater applicability to understanding human exposures to mutagenic substances and may provide more informative data for human cancer risk assessment.

## Acknowledgments

We thank Dr. Arun Pandiri and Ms. Julie Foley at the Division of Translational Toxicology for careful review of the manuscript and to members of TwinStrand who supported this work. Work performed for the National Toxicology Program, National Institutes of Environmental Health Sciences, National Institutes of Health, US Department of Health and Human Services was conducted under contract HHSN273201300009C (genetic toxicity testing) and ES103378-01. Work carried out at TwinStrand was supported by NIH R44 ES030642 to J.J.S.

## Data and Materials Availability

Sequencing data and metadata that support the findings of this study have been deposited in the NCBI Sequence Read Archive (SRA), accession number PRJNA932445.

