## Supplemental Data Figures and Tables for "Adopting Duplex Sequencing™ Technology for Genetic Toxicity Testing: A Proof-of-Concept Mutagenesis Experiment with N-Ethyl-N-Nitrosourea (ENU)-Exposed Rats"

### Supplementary Data Figure 1

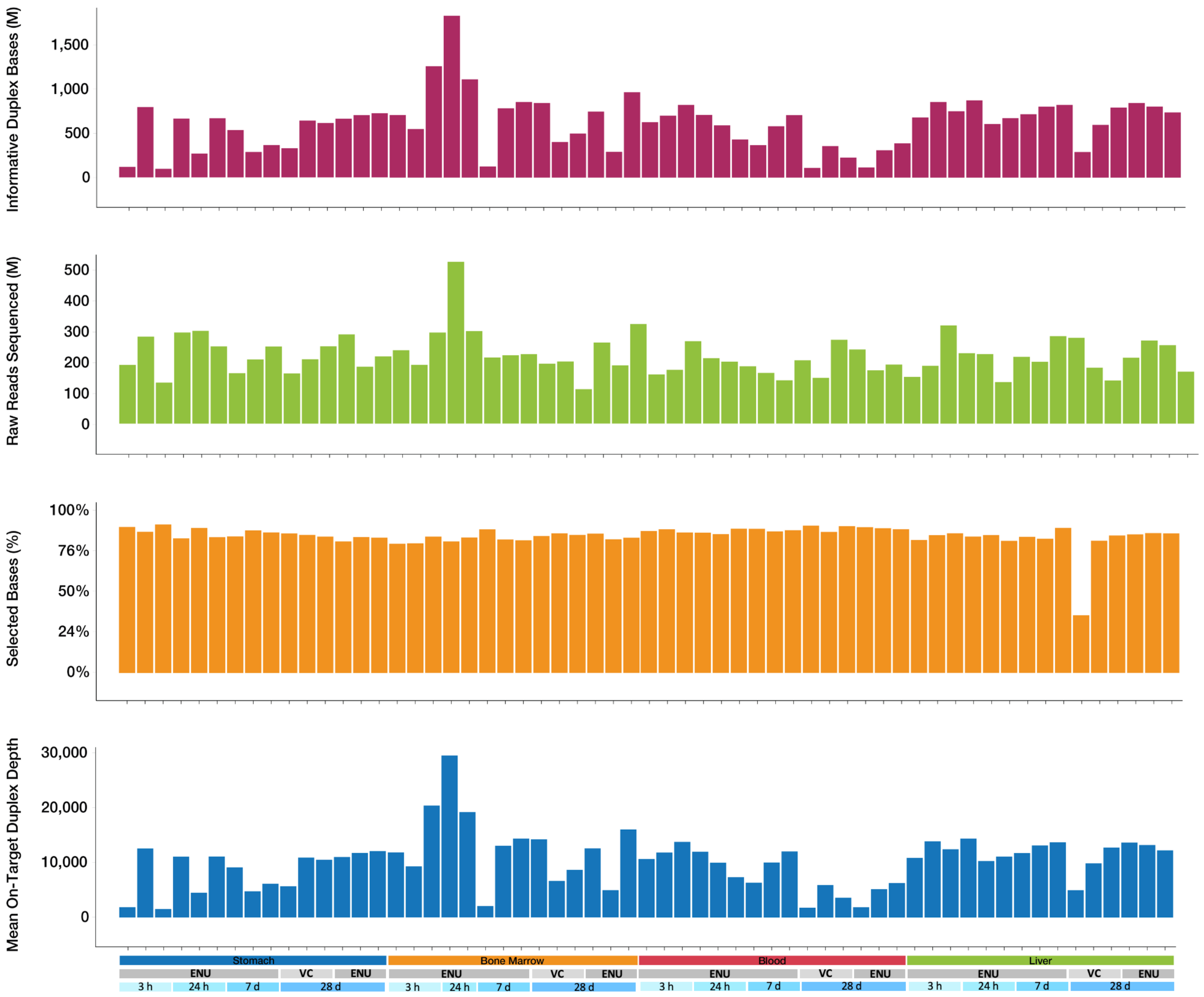


SD Fig. 1: The number of informative duplex bases per sample was variable due to differences in DNA quality and quantity. Three stomach samples, two from ENU-treated rats at the 3 h time point and one from the 24 h time point, had a low number of informative duplexes, possibly under-powering the measurement of SNVs.

### Supplementary Data Figure 2

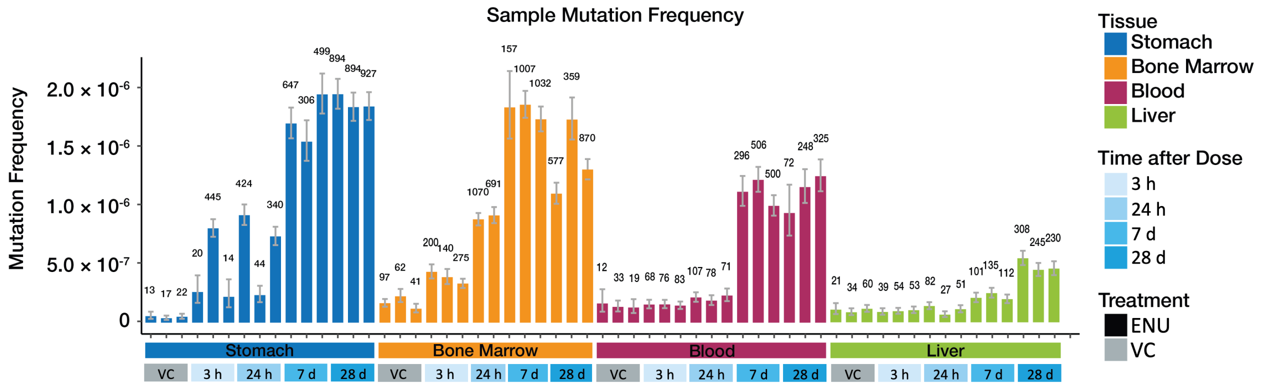


SD Fig. 2: Samples exhibited very low variability in MF among biological replicates except for stomach tissue at the 3 h and 24 h time points. The number of mutant bases detected per sample is noted above each bar. Error bars indicate the 95% confidence interval.

### Supplementary Data Figure 3

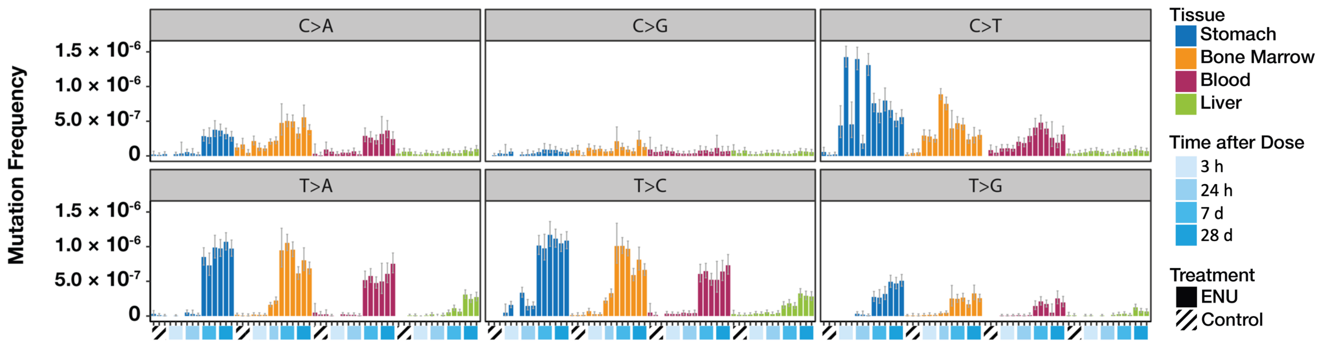


SD Fig. 3: Mutation frequency for each base substitution mutation subtype for each sample. Error bars represent 95% confidence interval.

### Supplementary Data Figure 4

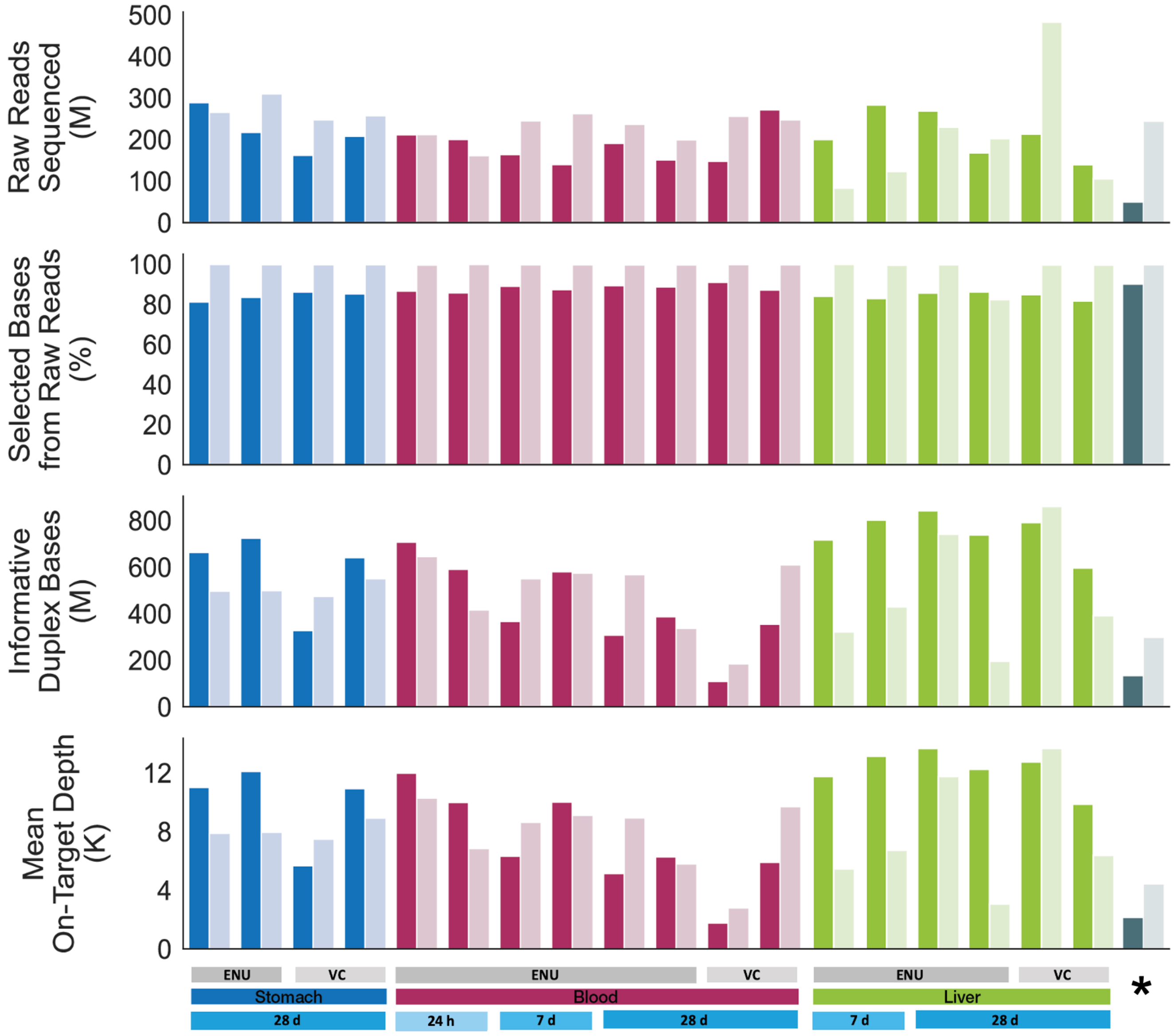


SD Fig. 4: Comparison of quality control indicators for duplex sequencing data obtained by TwinStrand versus ILS for a subset of samples. *Dark and light gray bars indicate TwinStrand versus DTT/ILS results for the TwinStrand kit's DNA control, respectively.

### Supplementary Data Figure 5

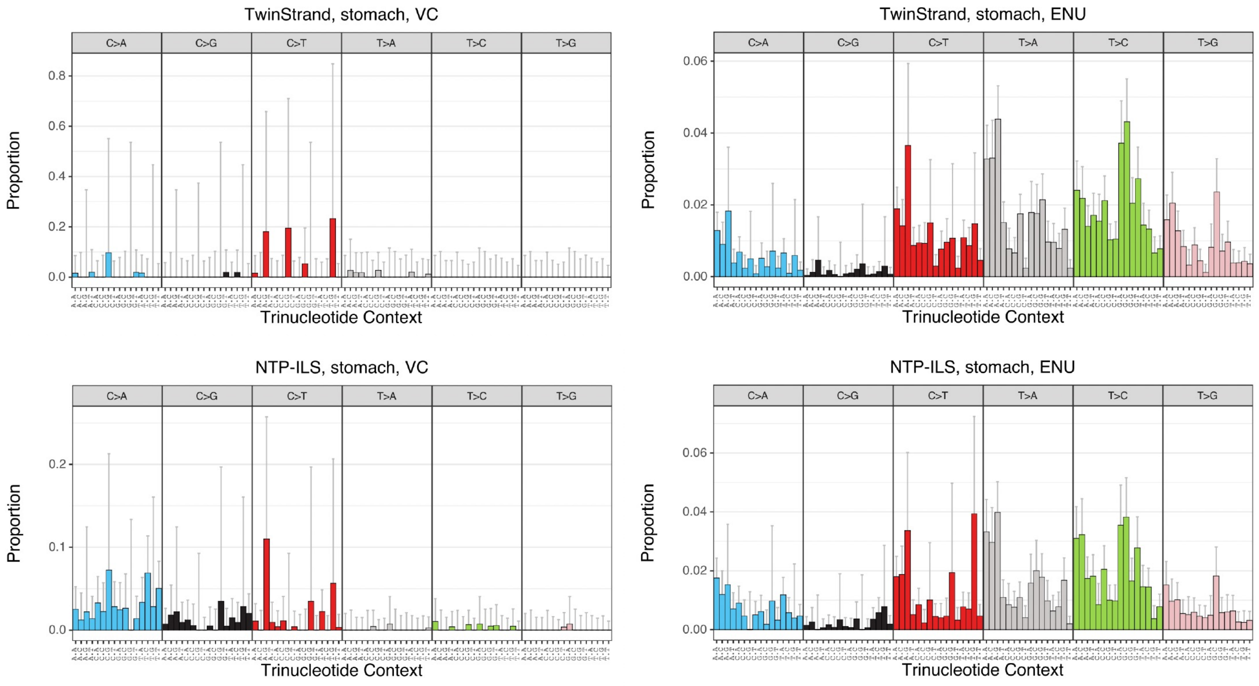


SD Fig. 5: Comparison of trinucleotide spectra obtained by TwinStrand versus ILS from a subset of stomach tissue samples from vehicle control (VC) or ENU-exposed rats.

### Supplementary Data Figure 6

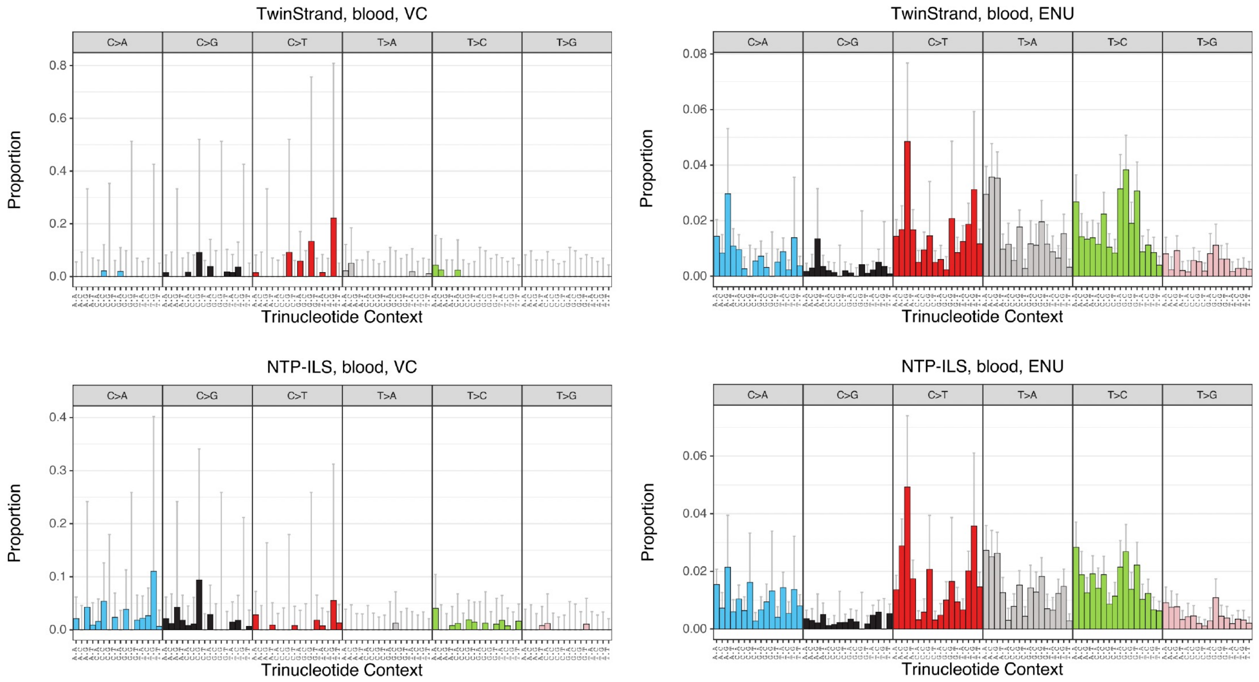


SD Fig. 6: Comparison of trinucleotide spectra obtained by TwinStrand versus ILS from a subset of blood samples from vehicle control (VC) or ENU-exposed rats.

### Supplementary Data Figure 7

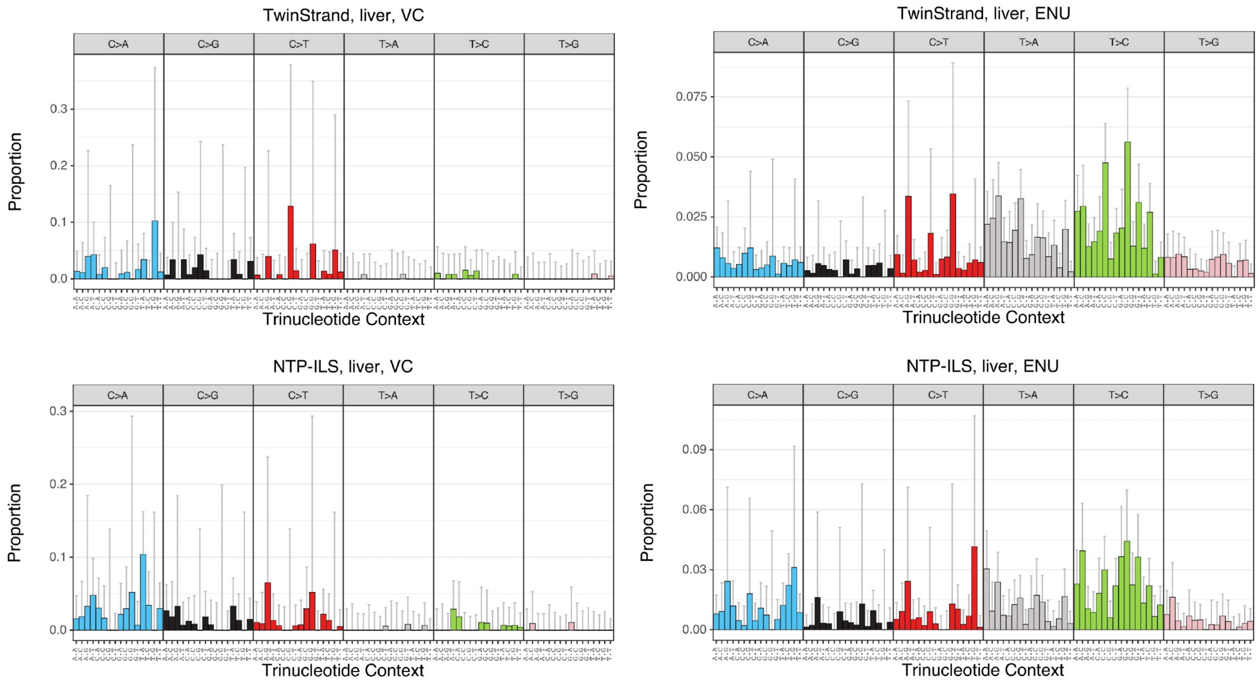


SD Fig. 7: Comparison of trinucleotide spectra obtained by TwinStrand versus ILS from a subset of liver tissue samples from vehicle control (VC) or ENU-exposed rats.

### Supplementary Data Figure 8

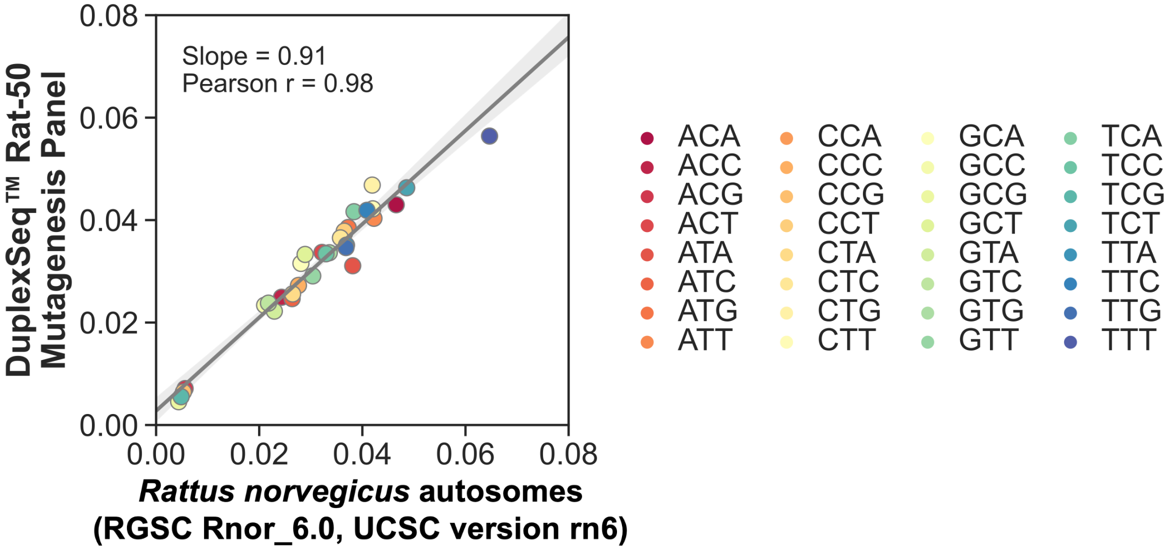


SD Fig. 8: The proportion of each pyrimidine-normalized trinucleotide sequence in the rat mutagenesis panel versus the rat autosome. The composition of trinucleotides in the panel and genome are proportional, ensuring unbiased 3-mer sampling when running the DuplexSeq$^{TM}$ Rat-50 Mutagenesis Assay.

### Supplementary Data Table 1

Supplementary Data Table 1: Analysis of contamination by DNA from non-rat species

| Sample Name (Animal ID) | Duplex consensus sequences classified as DNA from *Homo sapiens* | Total number of duplex consensus sequences | Fraction of duplex consensus classified as DNA from *Homo sapiens* |
|---|---|---|---|
| ENU 3 h bone marrow (2) | 1 | 3,776,900 | $2.65\times10^{-7}$ |
| ENU 3 h blood (2) | 1 | 5,122,180 | $1.95\times10^{-7}$ |
| ENU 24 h stomach (5) | 1 | 2,036,508 | $4.91\times10^{-7}$ |
| ENU 7 d blood (8) | 2 | 4,269,772 | $4.68\times10^{-7}$ |
| ENU 28 d bone marrow (15) | 1 | 2,185,114 | $4.58\times10^{-7}$ |
| ENU 28 d stomach (15) | 2 | 5,149,618 | $3.88\times10^{-7}$ |

### Supplementary Data Table 2

Supplementary Data Table 2: Mutation frequency for stomach tissue samples

| Exposure | Time Point | Animal ID | Mutations | Duplex Bases | Mutation Frequency | Mean | Lower CI[a] | Upper CI[a] |
|---|---|---|---|---|---|---|---|---|
| ENU | 3 h | 1 | 20 | 7,6371,874 | $2.62\times10^{-7}$ | | | |
| | | 2 | 445 | 552,103,920 | $8.06\times10^{-7}$ | $4.30\times10^{-7}$ | $-3.8\times10^{-7}$ | $12.4\times10^{-7}$ |
| | | 3 | 14 | 63,442,923 | $2.21\times10^{-7}$ | | | |
| ENU | 24 h | 4 | 424 | 461,373,568 | $9.19\times10^{-7}$ | | | |
| | | 5 | 44 | 187,171,246 | $2.35\times10^{-7}$ | $6.30\times10^{-7}$ | $-2.5\times10^{-7}$ | $15.1\times10^{-7}$ |
| | | 6 | 340 | 461,121,322 | $7.37\times10^{-7}$ | | | |
| ENU | 7 d | 7 | 647 | 379,886,484 | $17.0\times10^{-7}$ | | | |
| | | 8 | 306 | 197,740,046 | $15.5\times10^{-7}$ | $17.34\times10^{-7}$ | $12.3\times10^{-7}$ | $22.4\times10^{-7}$ |
| | | 9 | 499 | 255,625,934 | $19.5\times10^{-7}$ | | | |
| ENU | 28 d | 14 | 895 | 457,710,268 | $19.6\times10^{-7}$ | | | |
| | | 15 | 894 | 485,108,913 | $18.4\times10^{-7}$ | $18.83\times10^{-7}$ | $17.2\times10^{-7}$ | $20.5\times10^{-7}$ |
| | | 16 | 927 | 501,529,267 | $18.5\times10^{-7}$ | | | |
| Vehicle Control | 28 d | 10 | 13 | 237,403,015 | $0.55\times10^{-7}$ | | | |
| | | 11 | 17 | 454,906,602 | $0.37\times10^{-7}$ | $0.47\times10^{-7}$ | $0.25\times10^{-7}$ | $0.70\times10^{-7}$ |
| | | 12 | 22 | 439,557,944 | $0.50\times10^{-7}$ | | | |

[a]95% Wilson confidence intervals

### Supplementary Data Table 3

Supplementary Data Table 3: Mutation frequency for bone marrow tissue samples

| Exposure | Time Point | Animal ID | Mutations | Duplex Bases | Mutation Frequency | Mean | Lower CI[a] | Upper CI[a] |
|---|---|---|---|---|---|---|---|---|
| ENU | 3 h | 1 | 200 | 460,538,703 | $4.34\times10^{-7}$ | | | |
| | | 2 | 140 | 359,944,067 | $3.89\times10^{-7}$ | $3.86\times10^{-7}$ | $2.6\times10^{-7}$ | $5.1\times10^{-7}$ |
| | | 3 | 275 | 823,978,455 | $3.34\times10^{-7}$ | | | |
| ENU | 24 h | 4 | 1070 | 1,211,534,270 | $8.83\times10^{-7}$ | | | |
| | | 5 | ND[b] | ND | ND | $9.0\times10^{-7}$ | $6.8\times10^{-7}$ | $11.2\times10^{-7}$ |
| | | 6 | 691 | 753,434,243 | $9.17\times10^{-7}$ | | | |
| ENU | 7 d | 7 | 157 | 85,301,409 | $18.4\times10^{-7}$ | | | |
| | | 8 | 1007 | 540,323,967 | $18.6\times10^{-7}$ | $18.13\times10^{-7}$ | $16.5\times10^{-7}$ | $19.7\times10^{-7}$ |
| | | 9 | 1032 | 593,416,495 | $17.4\times10^{-7}$ | | | |
| ENU | 28 d | 14 | 577 | 523,217,268 | $11.0\times10^{-7}$ | | | |
| | | 15 | 359 | 206,729,355 | $17.4\times10^{-7}$ | $13.83\times10^{-7}$ | $5.7\times10^{-7}$ | $21.9\times10^{-7}$ |
| | | 16 | 870 | 664,529,293 | $13.1\times10^{-7}$ | | | |
| Vehicle Control | 28 d | 10 | 97 | 584,386,172 | $1.66\times10^{-7}$ | | | |
| | | 11 | 62 | 275,767,808 | $2.25\times10^{-7}$ | $1.70\times10^{-7}$ | $3.0\times10^{-7}$ | $3.8\times10^{-7}$ |
| | | 12 | 41 | 345,552,722 | $1.19\times10^{-7}$ | | | |

[a]95% Wilson confidence intervals
[b]ND = not determined

### Supplementary Data Table 4

Supplementary Data Table 4: Mutation frequency for blood samples

| Exposure | Time Point | Animal ID | Mutations | Duplex Bases | Mutation Frequency | Mean | Lower CI[a] | Upper CI[a] |
|---|---|---|---|---|---|---|---|---|
| ENU | 3 h | 1 | 68 | 440,964,413 | $1.54\times10^{-7}$ | | | |
| | | 2 | 76 | 489,163,406 | $1.55\times10^{-7}$ | $1.52\times10^{-7}$ | $1.4\times10^{-7}$ | $1.7\times10^{-7}$ |
| | | 3 | 83 | 572,055,400 | $1.45\times10^{-7}$ | | | |
| ENU | 24 h | 4 | 107 | 497,221,612 | $2.15\times10^{-7}$ | | | |
| | | 5 | 78 | 415,087,054 | $1.88\times10^{-7}$ | $2.12\times10^{-7}$ | $1.6\times10^{-7}$ | $2.7\times10^{-7}$ |
| | | 6 | 71 | 306,613,710 | $2.32\times10^{-7}$ | | | |
| ENU | 7 d | 7 | 296 | 264,461,052 | $11.2\times10^{-7}$ | | | |
| | | 8 | 506 | 414,450,099 | $12.2\times10^{-7}$ | $11.13\times10^{-7}$ | $8.4\times10^{-7}$ | $13.9\times10^{-7}$ |
| | | 9 | 500 | 500,749,035 | $10.0\times10^{-7}$ | | | |
| ENU | 28 d | 14 | 72 | 76,845,010 | $9.37\times10^{-7}$ | | | |
| | | 15 | 248 | 213,985,153 | $11.6\times10^{-7}$ | $11.16\times10^{-7}$ | $7.2\times10^{-7}$ | $15.2\times10^{-7}$ |
| | | 16 | 325 | 259,450,816 | $12.5\times10^{-7}$ | | | |
| Vehicle Control | 28 d | 10 | 12 | 73,393,847 | $1.64\times10^{-7}$ | | | |
| | | 11 | 33 | 245,485,418 | $1.34\times10^{-7}$ | $1.42\times10^{-7}$ | $0.95\times10^{-7}$ | $1.9\times10^{-7}$ |
| | | 12 | 19 | 147,519,258 | $1.29\times10^{-7}$ | | | |

[a]95% Wilson confidence intervals

### Supplementary Data Table 5

Supplementary Data Table 5: Mutation frequency for liver tissue samples

| Exposure | Time Point | Animal ID | Mutations | Duplex Bases | Mutation Frequency | Mean | Lower CI[a] | Upper CI[a] |
|---|---|---|---|---|---|---|---|---|
| ENU | 3 h | 1 | 39 | 443,727,392 | $0.88\times10^{-7}$ | | | |
| | | 2 | 54 | 573,421,255 | $0.94\times10^{-7}$ | $0.95\times10^{-7}$ | $0.76\times10^{-7}$ | $1.1\times10^{-7}$ |
| | | 3 | 53 | 515,597,719 | $0.10\times10^{-7}$ | | | |
| ENU | 24 h | 4 | 82 | 592,961,335 | $1.38\times10^{-7}$ | | | |
| | | 5 | 27 | 418,907,440 | $0.65\times10^{-7}$ | $1.05\times10^{-7}$ | $1.2\times10^{-7}$ | $2.0\times10^{-7}$ |
| | | 6 | 51 | 459,862,014 | $1.11\times10^{-7}$ | | | |
| ENU | 7 d | 7 | 101 | 484,313,896 | $2.09\times10^{-7}$ | | | |
| | | 8 | 135 | 543,250,415 | $2.49\times10^{-7}$ | $2.18\times10^{-7}$ | $1.5\times10^{-7}$ | $2.9\times10^{-7}$ |
| | | 9 | 112 | 569,196,793 | $1.97\times10^{-7}$ | | | |
| ENU | 28 d | 14 | 308 | 563,300,698 | $5.47\times10^{-7}$ | | | |
| | | 15 | 245 | 547,222,069 | $4.48\times10^{-7}$ | $4.84\times10^{-7}$ | $3.5\times10^{-7}$ | $6.2\times10^{-7}$ |
| | | 16 | 230 | 501,286,680 | $4.59\times10^{-7}$ | | | |
| Vehicle Control | 28 d | 10 | 21 | 196,508,264 | $1.07\times10^{-7}$ | | | |
| | | 11 | 34 | 399,311,323 | $0.85\times10^{-7}$ | $1.02\times10^{-7}$ | $0.64\times10^{-7}$ | $1.4\times10^{-7}$ |
| | | 12 | 60 | 522,612,706 | $1.15\times10^{-7}$ | | | |

[a]95% Wilson confidence intervals

### Supplementary Data Table 6

Supplementary Data Table 6: Mutation frequency variation analysis between biological replicates.

| Tissue | Exposure | Timepoint | Sample Size | Mean | Standard Deviation | Coefficient of Variation |
|---|---|---|---|---|---|---|
| Stomach | ENU | 3 h | 3 | $4.30\times10^{-7}$ | $3.27\times10^{-7}$ | 76.06% |
| | | 24 h | 3 | $6.30\times10^{-7}$ | $3.54\times10^{-7}$ | 56.19% |
| | | 7 d | 3 | $17.34\times10^{-7}$ | $2.04\times10^{-7}$ | 11.77% |
| | | 28 d | 3 | $18.83\times10^{-7}$ | $0.63\times10^{-7}$ | 3.37% |
| | VC | 28 d | 3 | $0.47\times10^{-7}$ | $0.09\times10^{-7}$ | 18.98% |
| Bone Marrow | ENU | 3 h | 3 | $3.86\times10^{-7}$ | $0.50\times10^{-7}$ | 13.05% |
| | | 24 h | 2 | $9.00\times10^{-7}$ | $0.24\times10^{-7}$ | 2.67% |
| | | 7 d | 3 | $18.13\times10^{-7}$ | $0.66\times10^{-7}$ | 3.65% |
| | | 28 d | 3 | $13.83\times10^{-7}$ | $3.23\times10^{-7}$ | 23.38% |
| | VC | 28 d | 3 | $1.70\times10^{-7}$ | $0.53\times10^{-7}$ | 31.32% |
| Blood | ENU | 3 h | 3 | $1.52\times10^{-7}$ | $0.06\times10^{-7}$ | 3.71% |
| | | 24 h | 3 | $2.12\times10^{-7}$ | $0.22\times10^{-7}$ | 10.42% |
| | | 7 d | 3 | $11.13\times10^{-7}$ | $1.11\times10^{-7}$ | 10.00% |
| | | 28 d | 3 | $11.16\times10^{-7}$ | $1.62\times10^{-7}$ | 14.53% |
| | VC | 28 d | 3 | $1.42\times10^{-7}$ | $0.19\times10^{-7}$ | 13.09% |
| Liver | ENU | 3 h | 3 | $0.95\times10^{-7}$ | $0.07\times10^{-7}$ | 7.88% |
| | | 24 h | 3 | $1.05\times10^{-7}$ | $0.37\times10^{-7}$ | 35.70% |
| | | 7 d | 3 | $2.18\times10^{-7}$ | $0.27\times10^{-7}$ | 12.44% |
| | | 28 d | 3 | $4.84\times10^{-7}$ | $0.54\times10^{-7}$ | 11.20% |
| | VC | 28 d | 3 | $1.02\times10^{-7}$ | $0.15\times10^{-7}$ | 15.01% |

### Supplementary Data Table 7

Supplementary Data Table 7: Comparison of mutation frequency (×$10^{-7}$) obtained by TwinStrand Biosciences, Inc. versus ILS, LLC for a subset of samples.

| Exposure | Animal ID | Tissue | Time Point | TwinStrand Biosciences, Inc. Mutation Frequency | TwinStrand Biosciences, Inc. Lower and Upper 95% CI | ILS, LLC Mutation Frequency | ILS, LLC Lower and Upper 95% CI |
|---|---|---|---|---|---|---|---|
| Vehicle Control | 10 | Blood | 28 d | 1.64 | 0.94, 2.86 | 1.55 | 0.98, 2.45 |
| | 11 | Blood | 28 d | 1.34 | 0.96, 1.89 | 1.78 | 1.42, 2.24 |
| | 11 | Liver | 28 d | 0.85 | 0.61, 1.19 | 1.64 | 1.22, 2.21 |
| | 12 | Liver | 28 d | 1.15 | 0.89, 1.48 | 1.42 | 1.14, 1.76 |
| | 10 | Stomach | 28 d | 0.55 | 0.32, 0.94 | 2.13 | 1.68, 2.70 |
| | 11 | Stomach | 28 d | 0.37 | 0.23, 0.60 | 2.33 | 1.89, 2.87 |
| ENU | 4 | Blood | 24 h | 2.15 | 1.78, 2.60 | 2.58 | 2.15, 3.11 |
| | 6 | Blood | 24 h | 1.88 | 1.51, 2.34 | 4.43 | 3.71, 5.28 |
| | 7 | Blood | 7 d | 11.2 | 10.0, 12.5 | 10.2 | 9.22, 11.3 |
| | 8 | Blood | 7 d | 12.2 | 11.2, 13.3 | 13.3 | 12.2, 14.5 |
| | 15 | Blood | 28 d | 11.6 | 10.2, 13.1 | 12.6 | 11.5, 13.8 |
| | 16 | Blood | 28 d | 12.5 | 11.2, 14.0 | 11.3 | 10.1, 12.7 |
| | 7 | Liver | 7 d | 2.09 | 1.72, 2.53 | 1.92 | 1.43, 2.75 |
| | 8 | Liver | 7 d | 2.49 | 2.10, 2.94 | 4.48 | 3.76, 5.35 |
| | 14 | Liver | 28 d | 5.47 | 4.89, 6.11 | 5.38 | 4.77. 6.06 |
| | 16 | Liver | 28 d | 4.59 | 4.03, 5.22 | 5.13 | 4.02, 6.56 |
| | 14 | Stomach | 28 d | 19.6 | 18.3, 20.9 | 20.9 | 19.4, 22.5 |
| | 16 | Stomach | 28 d | 18.5 | 17.3, 19.7 | 19.5 | 18.0, 21.0 |
| TwinStrand Control DNA | | | | 3.68 | 2.60, 5.19 | 3.21 | 2.49, 4.14 |
